# Impact of the radiated brain microenvironment on a panel of human patient-derived xenografts

**DOI:** 10.1101/2020.06.03.132365

**Authors:** Jibo Zhang, Ian E. Olson, Lucas P. Carlstrom, Masum Rahman, Karishma Rajani, Kshama Gupta, Libo Liu, Zhi Tang, Eliot F. Sananikone, Anqin (Vicky) Dong, Arthur E. Warrington, Moses Rodriguez, Jincao Chen, Mark A. Schroeder, Samar Ikram, Jann N. Sarkaria, Sandeep Burma, Terry C. Burns

## Abstract

**Objective:** Radiotherapy, combined with surgical resection and chemotherapy, remains a first-line treatment for infiltrative gliomas. However, these tumor are not surgically curable, and often recur, even within the prior radiation field, and may demonstrate a more aggressive phenotype. We recently demonstrated that the radiated brain tumor microenvironment promotes tumor aggressiveness in an orthotopic patient-derived xenograft (PDX) model of glioblastoma (Mayo GBM 143). Importantly, high grade gliomas display diverse molecular phenotypes, and whether this genetic variability leads to divergent behaviour in the radiated tumor microenvironment is unknown. Herein, we characterize the effects of the irradiated brain microenvinroment on nine additional unique GBM cell lines to better understand the nuances of how tumor molecular phenotypes influence cellular dynamics.

**Methods:** Female athymic nude mice were randomly divided into cranial radiation (15 Gy) and non-radiated groups. Mice then underwent intracranial implantation with one of the selected PDX GBM cell lines (GBM 6, 10, 12, 39, 46, 76, 123, 164, 196; total n=8-15, per group, per line). GBM 6 cells were additionally implanted 6 months after completion of fractionated radiation (4Gy × 10 fractions or 2Gy × 30 fractions) vs sham radiation. Kaplan-Meyer (K-M) and log-rank tests were performed to compare the survival between irradiated and non-irradiated groups.

**Result:** Of nine previously untested human GBM lines, we found that five demonstrated shorter survival in the pre-radiated brain (GBM 6, 46, 76, 164, 196); similar to previous observations with GBM 143. GBM 6 was also evaluated 6 months after fractionated radiation yielding similar results. However, two lines yielded prolonged survival in the pre-radiated brain (GBM 10, 12); GBM12 and 10 demonstrated the fastest baseline growth in the non-radiated brain; GBM 39, 123 whose rate of growth was not impacted by the radiated brain, demonstrated a an intermediate baseline growth rate between that of those positively and negatively impacted by the radiated brain microenvironment. No other clinical or molecular phenotype was found to consistently correlate with response to the radiated microenvironment.

**Conclusion:** Among a total of 10 total human GBM lines evaluated to date, 60% induce faster mortality in a radiated microenvironment, and 20% induce slower mortality. These results highlight the likely critical impact of the irradiated microenvironment on tumor behaviour, yet illustrate that different tumors may exhibit opposing responses. Although further evaluation will be needed to understand mechanisms of divergent behavior, our data suggest the increased rate of growth in the radiated microenvironment may not apply to the fastest-growing tumor lines, which could instead demonstrate a paradoxical response.

## Introduction

Glioblastoma (GBM) is the most common and deadly primary neoplasm of the adult central nervous system (CNS), with a median survival of 12-15 months following diagnosis [1]. Radiotherapy, combined with surgical resection and chemotherapy, remains a first-line treatment for GBM, but tumor recurrence is near absolute. These recurrences typically occur within the prior radiation field, demonstrating highly aggressive behavior and subsequence resistance to salvage therapy [23]. Unfortunately, when patients invariably experience tumor recurrence there are few clinical interventions that provide further survival benefit. As such, there remains an important unmet clinical need to identify factors that may contribute to early GBM recurrence despite maximal medical and surgical therapy.

Patient-derived xenograft (PDX) models of GBM are routinely employed to evaluate complex interactions between the human tumor cells and a CNS microenvironment [4,5,6]. These PDX models also serve as an invaluable preclinical model to evaluate the efficacy of candidate therapeutics [7,8]. The Mayo Clinic Glioma PDX National Resource comprises 96 well-characterized GBM lines that collectively capture much of the clinical and molecular breadth of human GBM [9].

A growing body of literature has elucidated the substantial impacts of radiotherapy on non-tumor cells present in the tumor microenvironment and surrounding CNS parenchyma [10,11]. Although radiation remains standard of care and despite the nearly uniform GBM recurrence rates, we and others have made the concerning observation that the radiated microenvironment actually increases the aggressiveness of GBM cells implanted into an irradiated brain [12,13,14]. To date, studies reporting this phenomena have each evaluated a single cell line, leaving open the question of how widespread this susceptibility to increased aggressiveness may be across the clinical and moleulular spectrum of human GBM. Indeed, all other studies to date besides our recent report in GBM143 have utilized rodent GBM cell lines. We thus sought to determine the behaviour of a representative cohort of additional GBM lines and thus selected nine additional GBM lines to encompass a diversity of clinical and molecular phenotypes spanning patient age, gender, recurrence, tumor growth rate, mutational phenotype, methylation status and molecular subtype (Table 1).

**Table 1.** Ten cell lines’ information

| Line | Sex | age | Recurrence Status | MGMT methylation | Subtype | IDH1 | CDKN2A | PTEN | EGFR | TP53 | Met | Tert Prom | Other |
| --- | --- | --- | --- | --- | --- | --- | --- | --- | --- | --- | --- | --- | --- |
| 6 | M | 65 | Primary | U | Clas |  | L | L | A (v3) | M | A | M |  |
| 10 | M | 41 | Recurrent | U | Mes |  | L | LM | A |  | A | M |  |
| 12 | M | 69 | Primary | M | Mes |  | L | L | AM | ML |  | M |  |
| 39 | M | 51 | Primary | M | Mes |  | L | LM | A (v3) |  | A | M | MDM4 & PIKC32B Amp |
| 46 | M | 56 | Recurrent | M | Mes |  | L | LM | A (v3) |  | A | M |  |
| 76 | M | 38 | Recurrent | M | Clas |  | L | LM | A (V3) |  | A | M |  |
| 123 | F | 62 | Primary | U | PN |  | L | L | A (V2) | M | A | M | CDK4&Myc Amp; ATRX mut |
| 143 | M | 68 | Secondary | U | Clas |  | L | L |  |  | A | M |  |
| 164 | F | 38 | Primary | M | PN | M<sup>h</sup> | L | L |  |  | A | - | PDGFR Amp; NF1 loss; ATRX trunc |
| 196 | F | 31 | Primary | M | ND | M<sup>hh</sup> | L | L | A | MA | A | - | PDGFR & RB1 Amp |

\* = dBT (maintained in culture). M=Mutant, A=Amplified, L=Loss
Subtypes: Classical (Clas), Mesenchymal (Mes), Proneural (P), Not determined (ND)
h heterozygous R132H
hh homozygous R132H

## Material and methods

### Ethics statement on mice

Studies were approved by the National Institutes of Health (NIH) and Mayo Clinic Institutional Animal Care and Use Committee (IACUC), Rochester, MN. All animal procedures were performed with proper animal handling, adhering to the NIH guidelines and protocols approved by the IACUC. Female athymic nude (heterozygous Hsd: Athymic Nude −Foxn1nu/Foxn1+ mice, aged 6–8 weeks from Charles River Laboratories, City, State) were randomly divided into irradiated and non-irradiated groups (4–10 mice per group; 8-15 mice total per group, per line). Mice were housed at the Mayo Clinic animal care facility, which is Association for Assessment and Accreditation of Laboratory and Animal Care International (AAALACI)-accredited.

### Cranial irradiation

Cranial irradiation was administered to isofluorane-anesthetic using the X-RAD SmART irradiator (Precision X-ray, North Branford, CT), which uses a cone beam CT (CBCT) for accurate target localization. The stereotactic coordinates were determined from the target-set on CBCT using the first scan for each mouse within all groups (values ranged between × = 0.25 to 0.35, y = −3.8 to −4.0, and z = −5.8 to −5.95, depending on mice and strain-type). Whole brain radiotherapy used parallel opposed lateral beams with 10mm square collimator. The radiation dose was 15Gy, single dose (Figure 1). Control group underwent identical management, except no radiation dose administered (0Gy). A separate cohort of mice additionally underwent fractionated radiation (2Gy × 30 fractions or 4Gy × 10 fractions, 5 days per week)

**Figure 1.**
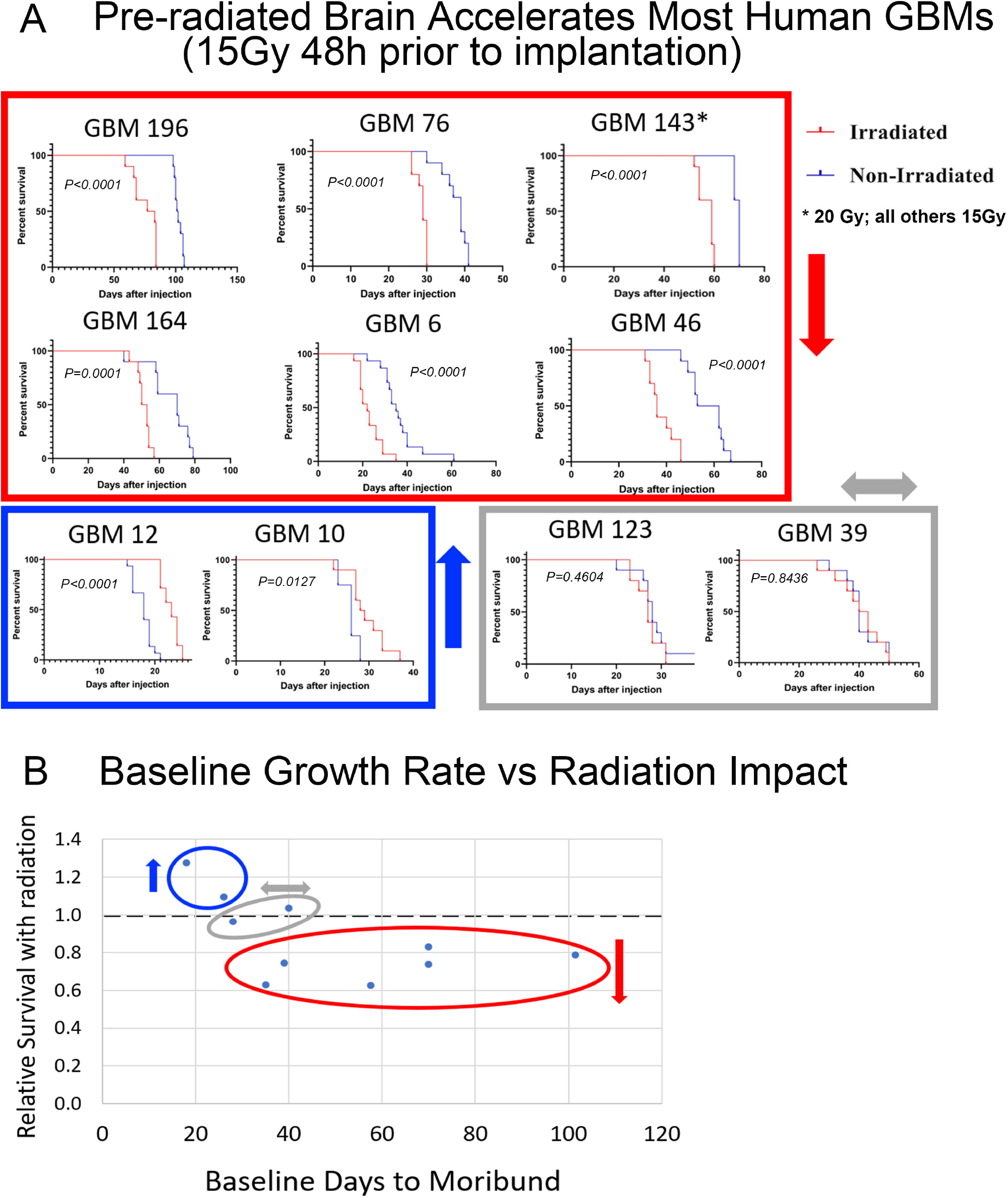
Experimental materials and methods.

### PDX GBM cell lines

A panel of well characterized PDX GBM cell lines (GBM 6, 10, 12, 39, 46, 76, 123, 143, 164, 196) were obtained from flank tumors and cultured in vitro (1-3 weeks). Detailed cell line information can be found in Table 1. Prior to implantation, cells were dissociated using TryplE (Cat# 12563011, Thermo Scientific) and resuspended in PBS at a concentration of 100,000 cells/µl, except for GBM 12 (33,333 cells/µl, given high baseline growth rate). Each mouse received 3uL of cell suspension.

### Intracranial injections

Intracranial injections (Figure 1) were performed as described by Carlson et al [15]. Mice were anesthetized using Ketamine: Xylazine mixture (100mg/kg Ketamine and 10mg/kg Xylazine, i.p.). After betadine prep and application of ophthalmic ointment (artificial tears), a 1cm midline incision exposed bregma, after which a burrhole was drilled 1mm anterior and 2mm lateral to bregma. Cells were implanted using a Kopf stereotactic frame and 26G Hamilton syringe a rate of 1µL/min for over 3min using a motorized injector. The needle was left in place for an additional 3min to minimize risk of reflux. Bone wax was applied to the burrhole, and the wound sutured with 4-0 vicryl. Triple antibiotic was applied to the incision and stitches to prevent infection, and the mouse was left in the warm cage to recover from anesthesia. Water was supplemented with children’s ibuprofen starting 48hrs prior to starting the procedure and continued for 48hrs post-surgery. The mice were monitored daily for a week. Sutures were removed 10 days post-surgery.

### Tissue Harvest and Processing

Mice were sacrificed at time of moribund, as described by Gupta SK, et al. [16]. Briefly, the sacrifice standards were as follows: hunched and lethargic; seizures; and weight loss of more than 20%. For sacrifice, deeply anestethized mice underwent transcranial perfusion with PBS followed by fromalin [17] prior to parrifin embedding and sectioning (Mayo Clinic Research Core Coordinator Animal Histology, Scottsdale, AZ). Histology was performed using haematoxylin and eosin (HE) stains. In some cohorts, a subset of brains were snap frozen and maintained for future studies.

### Statistical analysis

Kaplan-Meyer (K-M) curves were generated and analyzed using log-rank test to compare the survival between irradiated and non-irradiated groups with GraphPad Prism Version 8.2.1 (GraphPad Software Inc., San Diego, CA). A p-value less than 0.05 was considered statistically significant.

## Results

We evaluated the cellular phenotypes of nine GBM cell lines that were implanted into mice that received cranial radiation (15Gy) or sham treatment two days prior to tumor cell implantation (Table 2). Cell lines 6, 46, 76, 164, 196 demonstrated markedly reduced survival (21-40% shorter) when implanted into previously radiated animals (*P*<0.0005, each line). Animals implanted with GBM 10 or 12 experienced survival benefit in the irradiated mice (GBM 10: 27.8% prolonged survival, *P*<0.0001; GBM10: 9.6 prolonged survival; p=0.0127). Animals the recived injections with the remaining two lines (GBM 39, 123) demonstrated no difference in survival between the two groups(*P*>0.05) (Figure 2). Studies for GBM 6 and 12 were performed as initial and validation studies, yielding similar results (Supplemental data), and results pooled (Table 2).

**Table 2.** Mice number and median survival of all lines’ mice

| GBM | Mice Number |  | Median Survival (days) |  | *P* Value |
| --- | --- | --- | --- | --- | --- |
|  | Irradiated | Non-Irradiated | Irradiated | Non-Irradiated |  |
| **6** | 5 | 5 | 29 | 40 | 0.0050 |
| **6 repeated** | 10 | 10 | 19.5 | 32.5 | <0.0001 |
| **6 overalls** | 15 | 15 | 22 | 35 | <0.0001 |
| **10** | 10 | 8 | 28.5 | 26 | 0.0127 |
| **12** | 4 | 5 | 23 | 16 | 0.0069 |
| **12 repeated** | 10 | 10 | 23 | 19 | <0.0001 |
| **12 overalls** | 14 | 15 | 23 | 18 | <0.0001 |
| **39** | 10 | 10 | 41.5 | 40 | 0.8436 |
| **46** | 10 | 10 | 36 | 57.5 | <0.0001 |
| **76** | 10 | 10 | 29 | 39 | <0.0001 |
| **123** | 10 | 10 | 27 | 28 | 0.4604 |
| **143** | 10 | 10 | 58 | 70 | <0.0001 |
| **164** | 10 | 10 | 51.5 | 70 | 0.0001 |
| **196** | 10 | 10 | 80 | 101.5 | <0.0001 |

**Figure 2:**
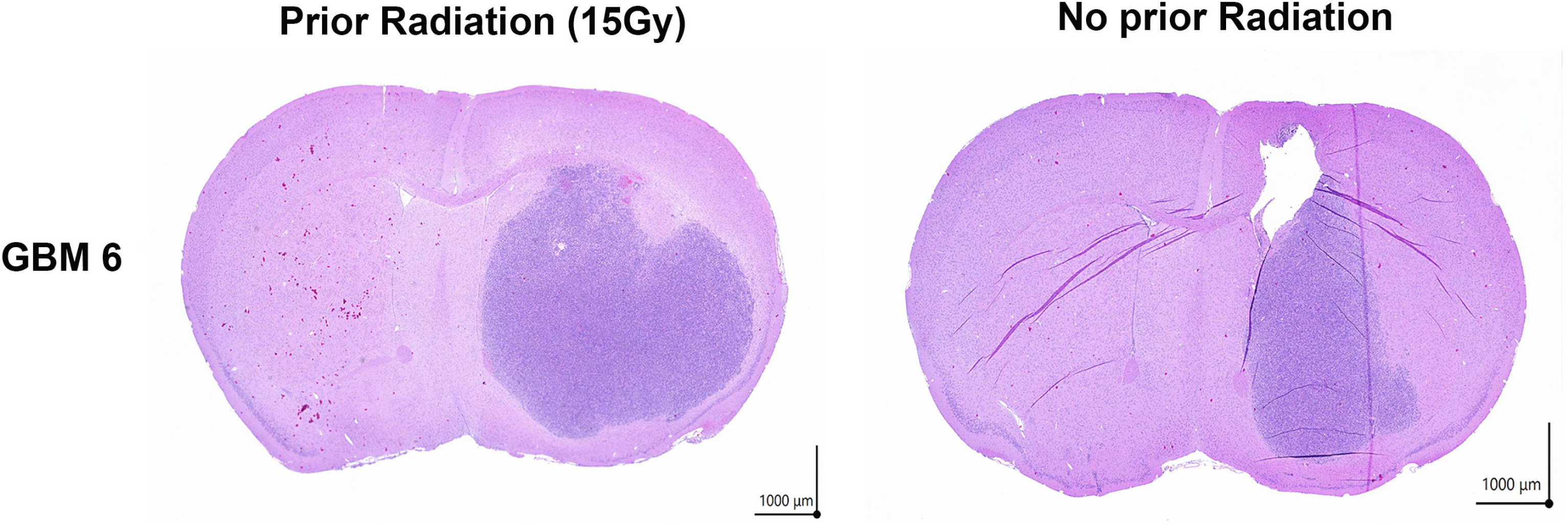
Impact of Prior Radiation on Human PDX growth rate. A: Kaplan Meier curves for 10 human PDX cell lines. 6, 46, 76, 143, 164, 196 (red box) demonstrated markedly reduced survival when implanted into previously radiated animals. Animals implanted with GBM 10 or 12 (Blue box) experienced survival benefit in the irradiated mice. Animals the received injections with GBM 39, 123 (grey box) demonstrated no difference in survival between the two groups. B: Baseline growth rate vs Radiation impact. Each cell line is plotted as a single data point. Cell lines in the blue, grey and red circles correspond to those in the blue grey and red boxes of Figure 2A.

Bioluminescent imaging (BLI) was performed over time in cohorts of mice implanted GBM cell lines 6 and 12 mice, which identified an accelerated tumor volume growth rate in irradiated GBM 6 mice, while GBM 12-implanted mice demonstrated a reduction from the normally very rapid rate of groth (Figure 3). A cohort of mice from GBM6 were sacficied at 21 days timepoint for future tissue analyses. A representative image showing typical tumor size in the pre-radiated vs non-radiated animals is shown in Figure 4.

**Figure 3:**
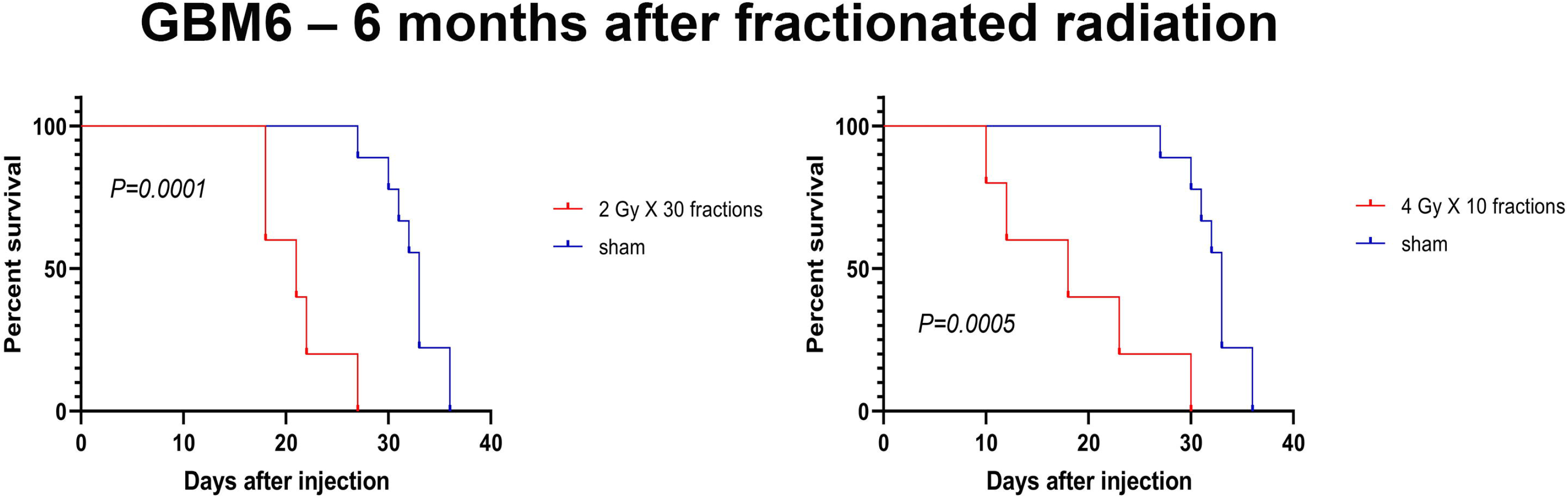
Opposite survival impacts of 15Gy pre-radiation 48h prior to implantation of GBM6 vs GBM12. A: Representative BLI images from mice implanted with GBM6 and 12 +/− 15Gy cranial radiation. B: BLI values on log scale during first 2 weeks of imaging. B: Overall growth curves until moribund on linear scale. * p<0.05; ** p<0.005. Error bars show standard deviation.

**Figure 4:**
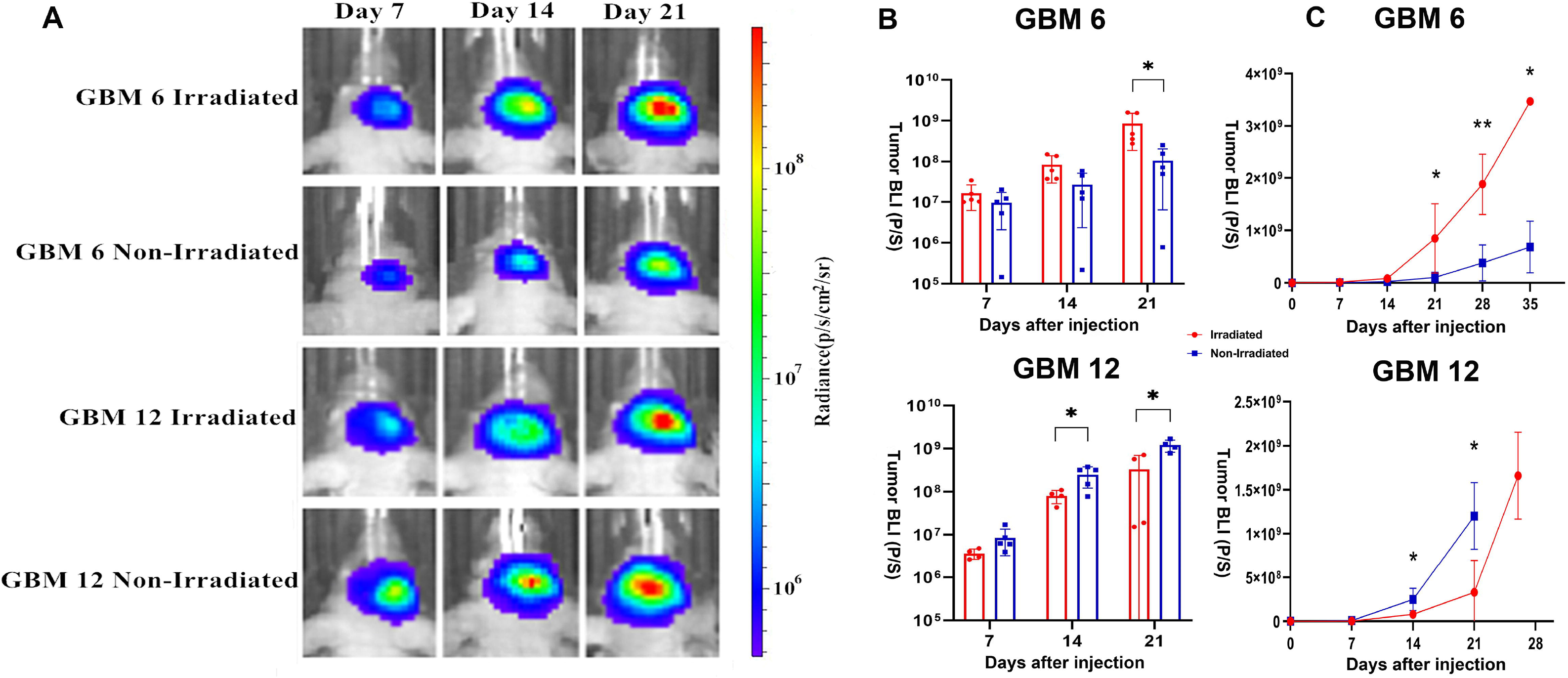
Tumor size in pre-radiated (48h) vs control mice sacrificed 21 days after implantation for GBM6. H&E of representative sections in representative mice shown. Findings are consistent with the significantly larger tumor size based on bioluminescence.

Since human patients with glioblastoma typically undergo fractionated radiation with expected recurrence several months after radiation, additional studies were performed using GBM 6 utilizing the identical 6 week fractionated radiation paradigm used for human patients (2Gy/d × 30 days, M-F for 6 weeks), as well as a hypofractionated regimen of 4Gy/d × 10 days M-F for 2 weeks. These clinically relevant fractionation doses yield similar biologically effective doses as 15Gy single fraction. Significantly accelerated mortality was observed in radiated as compared to sham mice for both fractionated radiation schemes (Figure 5).

**Figure 5:**
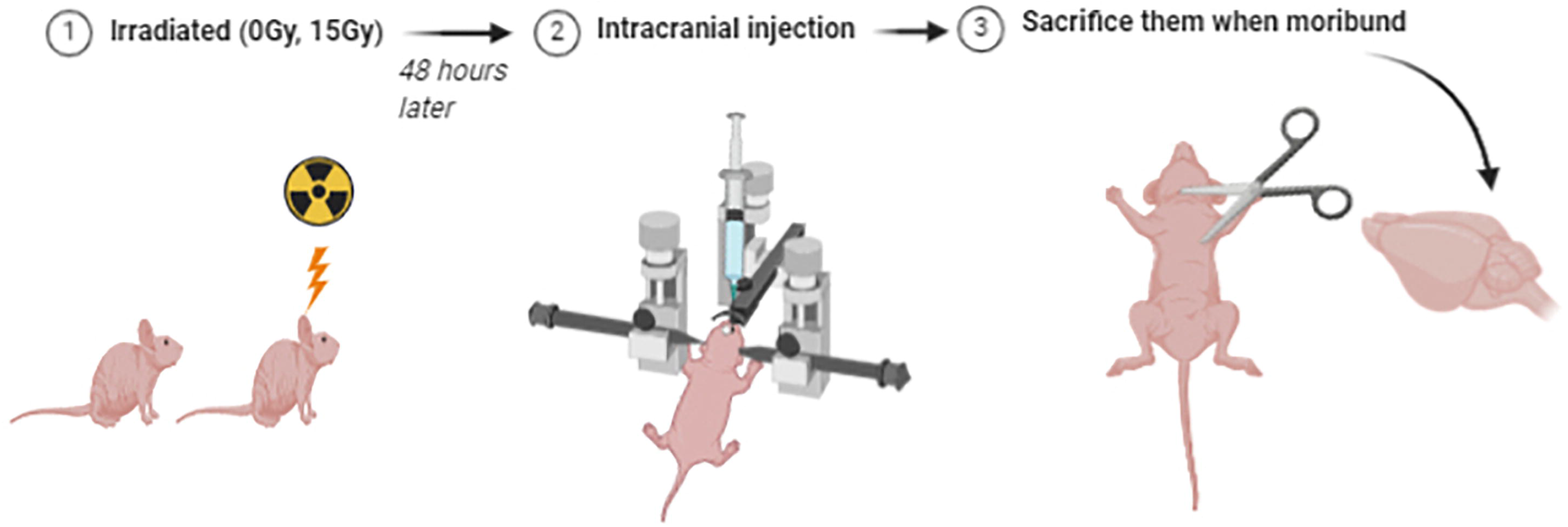
Worsened survival with fractionated radiation 6 months prior to tumor implantation. Kaplan Meier curves of animals implanted with GBM6 after undergoing 2 different regimens of fractionated radiation 6 months prior to implantation (4Gy × 10 fractions or 2Gy × 30 fractions). The same sham-radiated control group is demonstrated for comparison in each graph. No significant difference was observed in survival between animals undergoing 2Gy × 30 fractions vs 4Gy × 10 fractions.

## Discussion

These findings augment growing literature that illustrate the impact of the irradiated tumor microenvironment on tumor behavior [18]. Duan et al.[13] previously reported a radiation dose-dependent increased growth rate of mouse GBM cells within c57Bl6 mice upon delivery of (0, 20, 30, or 40 Gy by stereotactic radiation to the ipsilateral hemisphere. Desmarais and colleagues previously utilized the F98 rat model to demonstrate radiation-induced impacts on glioma behavior [19,20,21]. To our knowledge, this is the first effort to characterize the impact of a radiated brain microenvironment in multiple lines of human GBM.

Importantly, our findings suggest that the seemingly same tumor microenvironment may exert different impacts in a tumor-line-dependent manner, and may act to not only accelerate slower-growing lines, but also potentially slow the most aggressive cell lines under the experimental conditions used. Further work will be needed to determine the underlying mechanism of this phenomenon and to harness this knowledge to mitigate tumor aggressiveness. It is tempting to speculate that different factors may be responsible for the tumor-slowing impacts of the radiated microenvironment exerted upon GBM 10 and 12, as opposed to the tumor-accelerating factors observed in most lines, with the faster-growing cells exhibiting differential sensitivity to the signals directing slower growth. However, it is also conceivable that the same microenvironmental factor could exert divergent impacts in different cell lines.

Prior studies have demonstrated that CXCL12 (SDF1) [22], CSF1R [23] and IGF1 [24] in can facilitate tumor recurrence through vasculogenesis, macrophage polarization and stroma-mediated PI3K signaling, respectively. Hypoxia itself as may be induced by radiation has been shown to promote M2-like macrophage polarization [25]. We previously reported that brain radiation induces a unique microglial transcriptome that is somewhat more closerly related to M1 and M2, with strong aging-like features that could impact tumor aggressiveness [26]. To date, the impact of radiation upon other CNS cell types has been incompletely characterized, leaving open the possibility for complex interactions between the radiated brain and tumor. Indeed, the cognitive sequelae of MTX has been associated with altered physiology in astrocytes, microglia and oligodendrocyte progenitor cells [27], while neural activity itself has been shown to regulate glioma growth [28,29]. Identifying differential impacts of radiation on different GBM lines adds to the growing inventory of available tools to dissect molecular mechanisms of microenvironmental impacts on gioma behavior. Since Radiation is known to substantially impact microglia, OPCs, Myelin [30], endothelium [31], astrocytes [32], neurons [33], and macrophages [34], as well as the glioma itself, a dynamic milieu of signaling molecules including metabolites and cytokines may mediate interactions of fluctuating importance within a heterogeneously evolving tumor ecosystem [35].

In this study, ten well-characterized PDX GBM cell lines, providing a cross-sectional snapshot of human cellular responses to the radiated microenvironment. That a radiation dose response has also previously been shown to increase tumor growth in immunocompetent rodent GBM models suggests primary mechanisms other than adaptive immunity. Other host features that could impact results but not evaluated include age, gender, background strain, and implant location. For example, whether accelerated tumor growth would occur in the pre-radiated flank, rather than brain, is unknown. However, oncreased macrophage activation associated with a senesescent microenvironment has been shown to increase the infiltration of metastatic tumors [36]

At this point, multiple othe questions remain unanswered. Although we have demonstrated the reproducibility of the radiated microenvironment accelerating GBM6 for at least 6 months, and with multiple radiation paradigms, we currently have relatively little information about the growth-inhibiting characteristics observed in GBM10 and 12. Although these lines are both mesenchymal, so too is GBM46, which grew faster in the radiated brain. Further lines would need to be tested to determine if the apparent association between growth inhibition and baseline rapid growth is a generalizable phenomenon. Our prior study demonstrated that animals implanted with GBM143 at 2 months after radiation demonstrated significantly shorter survival than mice implanted with cells 2 days after radiation. Whether this temporal correlation between faster tumor growth and later timepoint after radiation is reproducible in other cell lines is unknown. If 11 true, results showing increasd growth after implantation at 48h could prove an underestimate of impacts at later post-radiation timepoints.

Of clinical relevance, our data suggest that perhaps a majority of patient’s GBM could recur in a miroevnironment that is more conducive to their growth at time of recurrence after radiation than was the case when the tumor first appeared and radiation had not occurred. Younger patients with typically lower grarde IDH-mutant tumors must must sometimes decide whether or not to delay radiation, with the expectation of needing further surgery at an earlier timepoint. Since 2 IDH-mutant gliomas (GBM164 and 196) both showed faster growth in the radiated microenvironment, these results—albeit limited by the murine model could lend support to delaying radiation when safe and feasible to do so. Whether similar impacts would be seen in a brain pre-treated with chemotherapy, and whether the tumor is any more or less sensitive to TMZ in the pre-radiated brain are remain open but clinically relevant questions. However, other lasting impacts of prior chemotherapy such as MTX on the brain microenvironment have been well established [37].

## Conclusion

In summary, we demonstrate that a majority of evaluated human GBM lines develop a more aggressive phenotype if implanted into the previously radiated brain—an effect that lasts at least 6 months after radiation. However, we also observed that implantion 48 after 15Gy radiation also served to attenuate the growth of the 2 fastest-growing cell lines tested (GBM10 and 12). It is hoped that further studies to understand the cellular and molecular mechanisms responsile for this enhanced or suppressed tumor volume growth could reveal opportunities to mitigate the aggressiveness of recurrent glioblastoma.

## Acknowlegements

Rehan Saber, Rujapope Sutiwisesak, Moustafa A. Mansour, Li Wang, Brytanny Howard

## Statement of Non-duplication

We certify that this manuscript is a unique submission and is not being considered for publication by any other source in any medium. Further, the manuscript has not been published, in part or in full, in any form.

## Ethical approval

Studies were approved by the National Institutes of Health (NIH) and Mayo Clinic Institutional Animal Care and Use Committee (IACUC), Rochester, MN. All animal procedures were performed with proper animal handling, adhering to the NIH guidelines and protocols approved by the IACUC.

## Authors’ contributions

Project conception: IEO, EFS, MRo, SB, TCB. Experimental design: IEO, JC, MAS, TCB.

Contributed cell lines, reagents or protocols: EFS, JNS, SB.

Performed experiments: JZ, IEO, MRa, KR, KG, LL, AD, AEW, MAS, SI Statistical analysis: JZ, IEO, KG, TCB.

Wrote the Manuscript: JZ, IEO, LPC, TCB.

## Availability of data and materials

The data that support the findings of this study are available on request from the corresponding author.

## References

1 Ostrom QT, Cioffi G, Gittleman H, et al. CBTRUS Statistical Report: Primary Brain and Other Central Nervous System Tumors Diagnosed in the United States in 2012-2016. Neuro Oncol. 2019;21(Suppl 5):v1–v100

2 Jeon Hee-Young, Kim Jun-Kyum, Ham Seok Won, et al. Irradiation induces glioblastoma cell senescence and senescence-associated secretory phenotype. Tumour Biol., 2016; 37: 5857–67.

3 Nizamutdinov D, Stock EM, Dandashi JA, et al. Prognostication of Survival Outcomes in Patients Diagnosed with Glioblastoma. World Neurosurg. 2018;109:e67–e74.

4 Yue Zhao, Timothy Wai Ho Shuen, Tan Boon Toh, et al. Development of a New Patient-Derived Xenograft Humanised Mouse Model to Study Human-Specific Tumour Microenvironment and Immunotherapy. Gut. 2018;67(10):1845–1854.

5 Jung J, Seol HS, Chang S. The Generation and Application of Patient-Derived Xenograft Model for Cancer Research. Cancer Res Treat. 2018;50(1):1–10.

6 Lai Y, Wei X, Lin S, et al. Current status and perspectives of patient-derived xenograft models in cancer research. J Hematol Oncol. 2017;10(1):106.

7 Yu D, Khan OF, Suvà ML, et al. Multiplexed RNAi therapy against brain tumor-initiating cells via lipopolymeric nanoparticle infusion delays glioblastoma progression. Proc Natl Acad Sci U S A. 2017;114(30):E6147–E6156.

8 Ramachandran M, Yu D, Dyczynski M, et al. Safe and Effective Treatment of Experimental Neuroblastoma and Glioblastoma Using Systemically Delivered Triple MicroRNA-Detargeted Oncolytic Semliki Forest Virus. Clin Cancer Res. 2017;23(6):1519–1530.

9 Vaubel RA, Tian S, Remonde D, et al. Genomic and Phenotypic Characterization of a Broad Panel of Patient-Derived Xenografts Reflects the Diversity of Glioblastoma. Clin Cancer Res. 2020;26(5):1094‐1104. 9

10 Nasrollah Jabbari, Muhammad Nawaz, Jafar Rezaie. Bystander Effects of Ionizing Radiation: Conditioned Media From X-ray Irradiated MCF-7 Cells Increases the Angiogenic Ability of Endothelial Cells. Cell Commun Signal. 2019;17(1):165.

11 Sundgren PC, Cao Y. Brain irradiation: effects on normal brain parenchyma and radiation injury. Neuroimaging Clin N Am. 2009;19(4):657–668.

12 Seo YS, Ko IO, Park H, et al. Radiation-Induced Changes in Tumor Vessels and Microenvironment Contribute to Therapeutic Resistance in Glioblastoma. Front Oncol. 2019;9:1259.

13 Duan C, Yang R, Yuan L, et al. Late Effects of Radiation Prime the Brain Microenvironment for Accelerated Tumor Growth. Int J Radiat Oncol Biol Phys. 2019;103(1):190–194.

14 Gupta K, Burns TC. Radiation-Induced Alterations in the Recurrent Glioblastoma Microenvironment: Therapeutic Implications. Front Oncol. 2018;8:503.

15 Carlson BL, Pokorny JL, Schroeder MA, Sarkaria JN. Establishment, maintenance and in vitro and in vivo applications of primary human glioblastoma multiforme (GBM) xenograft models for translational biology studies and drug discovery. Curr Protoc Pharmacol. 2011;Chapter 14:Unit 14.16.

16 Gupta SK, Kizilbash SH, Carlson BL, et al. Delineation of MGMT Hypermethylation as a Biomarker for Veliparib-Mediated Temozolomide-Sensitizing Therapy of Glioblastoma. J Natl Cancer Inst. 2015;108(5).

17 Gage GJ, Kipke DR, Shain W. Whole animal perfusion fixation for rodents. J Vis Exp. 2012;(65). pii: 3564.

18 Wild-Bode C, Weller M, Rimner A, et al. Sublethal irradiation promotes migration and invasiveness of glioma cells implications for radiotherapy of human glioblastoma. Cancer Res 2001;61:2744–2750.

19 Desmarais G, Fortin D, Bujold R, Wagner R, Mathieu D, Paquette B. Infiltration of glioma cells in brain parenchyma stimulated by radiation in the F98/Fischer rat model. Int J Radiat Biol. 2012;88(8):565–574.

20 Desmarais G, Charest G, Fortin D, Bujold R, Mathieu D, Paquette B. Cyclooxygenase-2 inhibitor prevents radiation-enhanced infiltration of F98 glioma cells in brain of Fischer rat. Int J Radiat Biol. 2015;91(8):624–633. 6

21 Desmarais G, Charest G, Therriault H, et al. Infiltration of F98 glioma cells in Fischer rat brain is temporary stimulated by radiation. Int J Radiat Biol. 2016;92(8):444–450.

22 Kioi M, Vogel H, Schultz G, Hoffman RM, Harsh GR, Brown JM. Inhibition of vasculogenesis, but not angiogenesis, prevents the recurrence of glioblastoma after irradiation in mice. J Clin Invest. 2010;120(3):694–705.

23 Yan D, Kowal J, Akkari L, et al. Inhibition of colony stimulating factor-1 receptor abrogates microenvironment-mediated therapeutic resistance in gliomas. Oncogene. 2017;36(43):6049–6058.

24 Quail DF, Bowman RL, Akkari L, et al. The tumor microenvironment underlies acquired resistance to CSF-1R inhibition in gliomas. Science. 2016;352(6288):aad3018.

25 Leblond MM, Gérault AN, Corroyer-Dulmont A, et al. Hypoxia induces macrophage polarization and re-education toward an M2 phenotype in U87 and U251 glioblastoma models. Oncoimmunology. 2015;5(1):e1056442.

26 Li MD, Burns TC, Kumar S, Morgan AA, Sloan SA, Palmer TD. Aging-like changes in the transcriptome of irradiated microglia. Glia. 2015;63(5):754–767.

27 Gibson EM, Nagaraja S, Ocampo A, et al. Methotrexate Chemotherapy Induces Persistent Tri-glial Dysregulation that Underlies Chemotherapy-Related Cognitive Impairment. Cell. 2019;176(1-2):43–55.e13.

28 Venkatesh HS, Johung TB, Caretti V, et al. Neuronal Activity Promotes Glioma Growth through Neuroligin-3 Secretion. Cell. 2015;161(4):803–816.

29 Venkatesh HS, Tam LT, Woo PJ, et al. Targeting neuronal activity-regulated neuroligin-3 dependency in high-grade glioma. Nature. 2017;549(7673):533–537.

30 Burns TC, Awad AJ, Li MD, Grant GA. Radiation-induced brain injury: low-hanging fruit for neuroregeneration. Neurosurg Focus. 2016;40(5):E3.

31 Lyubimova N, Hopewell JW. Experimental evidence to support the hypothesis that damage to vascular endothelium plays the primary role in the development of late radiation-induced CNS injury. Br J Radiol. 2004;77(918):488–492.

32 Turnquist C, Beck JA, Horikawa I, et al. Radiation-induced astrocyte senescence is rescued by ∆133p53. Neuro Oncol. 2019;21(4):474–485.

33 Wu PH, Coultrap S, Pinnix C, et al. Radiation induces acute alterations in neuronal function. PLoS One. 2012;7(5):e37677.

34 Leblond MM, Pérès EA, Helaine C, et al. M2 macrophages are more resistant than M1 macrophages following radiation therapy in the context of glioblastoma. Oncotarget. 2017;8(42):72597–72612.

35 Schäfer N, Gielen GH, Rauschenbach L, et al. Longitudinal heterogeneity in glioblastoma: moving targets in recurrent versus primary tumors. J Transl Med. 2019;17(1):96.

36 Coppé JP, Desprez PY, Krtolica A, Campisi J. The senescence-associated secretory phenotype: the dark side of tumor suppression. Annu Rev Pathol. 2010;5:99–118.

37 Geraghty AC, Gibson EM, Ghanem RA, et al. Loss of Adaptive Myelination Contributes to Methotrexate Chemotherapy-Related Cognitive Impairment. Neuron. 2019;103(2):250–265.e8.

