## Supplementary figures and images for "Impact of the radiated brain microenvironment on a panel of human patient-derived xenografts"

### Supplemental data

#### Supplemental Figure: Replication studies (GBM 6 and 12) 2 day after 15Gy

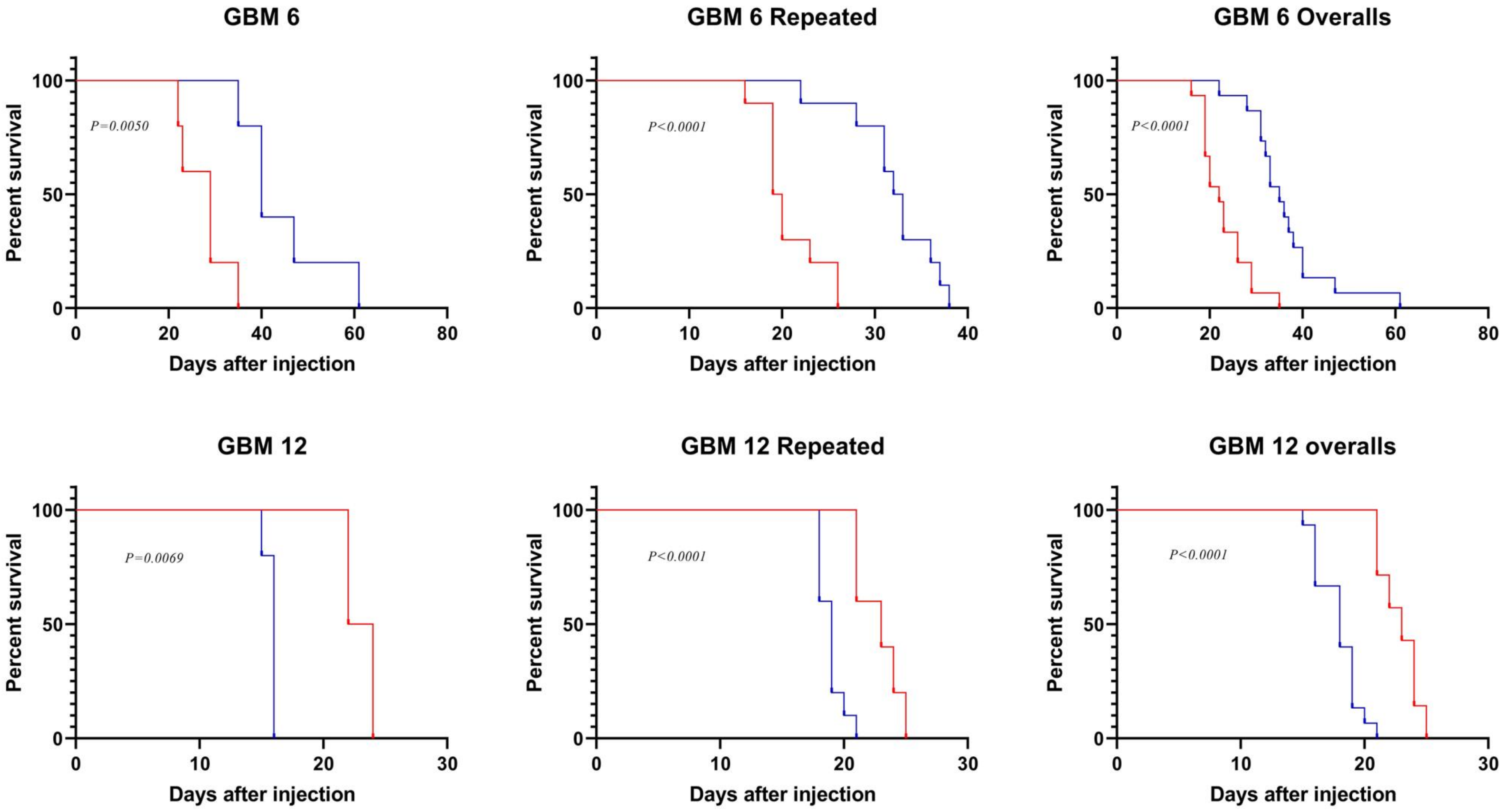
